# A comparison of telencephalon composition among chickens, junglefowl, and wild galliforms

**DOI:** 10.1101/2023.09.10.557096

**Authors:** Kelsey J. Racicot, Jackson R. Ham, Jacqueline Augustine, Rie Henriksen, Dominic Wright, Andrew N. Iwaniuk

## Abstract

Domestication is the process of modifying animals for human benefit through selective breeding in captivity. One of the traits that often diverges is the size of the brain and its constituent regions; almost all domesticated species have relatively smaller brains and brain regions than their wild ancestors. Although the effects of domestication on the brain have been investigated across a range of both mammal and bird species, almost nothing is known about the neuroanatomical effects of domestication on the world’s most common bird: the chicken (*Gallus gallus*). We compared the quantitative neuroanatomy of the telencephalon of white leghorn chickens with red junglefowl, their wild counterpart, and several wild galliform species. We focused specifically on the telencephalon because telencephalic regions typically exhibit the biggest differences in size in domesticate-wild comparisons. Relative telencephalon size was larger in chickens than in junglefowl and ruffed grouse (*Bonasa umbellus*). The relative size of telencephalic regions did not differ between chickens and junglefowl but did differ in comparison with ruffed grouse. Ruffed grouse had larger hyperpallia and smaller entopallial, nidopallial and striatal volumes than chickens and junglefowl. Multivariate analyses that included an additional three wild grouse species corroborated these findings: chicken and junglefowl have relatively larger nidopallial and striatal volumes than grouse. Conversely, the mesopallial and hyperpallial volumes tended to be relatively smaller in chickens and junglefowl. From this suite of comparisons, we conclude that chickens do not follow a pattern of widespread decreases in telencephalic region sizes that is often viewed as typical of domestication. Instead, chickens have undergone a mosaic of changes with some regions increasing and others decreasing in size and there are few differences between chickens and junglefowl.

## Introduction

Domestication of animals began approximately 15,000 years ago with wolves (*Canis lupus*), soon to be followed by many other plants and animals (DeMello, 2012; Vigne, 2011). The process of domesticating animals is almost identical across species: a subset of a wild population is isolated and then selected for specific traits within a captive environment across several generations. Some of the desirable traits that are actively selected for include tameness towards humans, larger body sizes (in meat producing animals), certain coat or plumage colors, and specific behavioural traits (Price, 1984; Trut, 1999). In addition to artificial selection, animals must also adapt to a human-made environment and, in most instances, different food sources than what they would normally eat in the wild (Price, 1984). Over time, artificial selection and captive breeding ultimately changes their phenotype such that they differ significantly in phenotype from the wild type (DeMello, 2012).

Domestication not only alters those traits under direct selection by humans, but there are also indirect effects. Brain size is frequently cited as a trait that changes as a result of domestication (Sanchez-Villagra 2022; Hecht et al. 2023); when domesticated mammals are compared with their wild counterparts, the domesticates (domesticated strains of a species) typically have relatively smaller brains (Kruska 2005). Similar patterns are observed in many domesticated bird species. For example, in ducks (*Anas platyrhyncos*) and pigeons (*Columba livia*), overall brain size decreases by 14% and 7%, respectively (Ebinger, 1995; Ebinger & Löhmer, 1984). Some of this reduction in relative brain size occurs as a result of larger body sizes in domesticates (Henriksen et al., 2016), but many telencephalic regions also become disproportionately smaller in domesticates. Indeed, the most drastic volumetric changes in the brain associated with domestication are decreases in the sizes of the sensory cortices, hippocampus, and amygdala (Brusini et al., 2018; Ebinger & Löhmer, 1984; Kruska, 1988). Changes in the sizes of brain regions are not, however, consistent across species and some regions do not change in size at all in domesticated birds. For example, the arcopallium is greatly reduced in volume in both domestic ducks (Ebinger 1995) and geese (*Anser anser*, Ebinger and Lohmer 1987), but not in domestic pigeons (Ebinger and Lohmer 1984). Similarly, hyperpallial regions are reduced in domestic pigeons and geese (Ebinger & Löhmer, 1984, 1987), but not in ducks (Ebinger, 1995). There are even examples of the expansion of telencephalic regions in domesticates, despite a reduction in overall brain size, such as the enlarged hippocampal formation of homing pigeons relative to wild rock doves (Rehkämper et al., 2008). Similarly, Henriksen et al. (2016) showed that white leghorn chickens (*Gallus gallus domesticus*) have relatively smaller brains than red junglefowl (*G. g. gallus*) but have proportionately larger cerebella and telencephala. Thus, brain region sizes in domesticated birds can be either larger or smaller than their ancestral counterparts.

Many domesticated species have been the focus of neuroanatomical studies, but data is lacking for the world’s most common bird species: the chicken. All domestic chicken breeds are derived from wild red junglefowl (*Gallus gallus gallus*), a species found in south-east Asia (Wang et al., 2020). Red junglefowl were first domesticated in 5000 B.C. and their original purpose was cockfighting for entertainment (Al-Nasser et al., 2007; Perry-Gal et al., 2015).

Three-thousand years after their original domestication, they had spread to the Middle East and Europe (Wang et al., 2020), at which point chickens had become a food source selected for both larger body size and egg production. Today, there are hundreds of chicken breeds that differ in size, behaviour, and morphology from the red junglefowl (Al-Nasser et al., 2007; Perry-Gal et al., 2015). Brain composition varies among many of these breeds, and some of that variation may reflect behaviour (Rehkämper et al. 2003). For example, the white crested Polish chicken has the most divergent brain of the breeds examined thus far, with an enlarged telencephalon, optic tract, and diencephalon compared with other breeds (Frahm & Rehkamper, 1998). Despite these broad comparisons across chicken breeds, neuroanatomical data on red junglefowl are lacking.

Compared to red junglefowl, domestic chickens are tamer towards humans and generally have larger bodies (Campler et al., 2009; Henriksen et al., 2016), but relatively smaller brains (Henriksen et al., 2016). As mentioned previously, chickens have a relatively larger telencephalon than junglefowl, but whether this is the result of coordinated increases across all telencephalic regions or expansion of only a few regions remains unknown. Determining which regions are relatively larger or smaller in chickens would provide new insights into behavioural differences between chickens and junglefowl (Schütz and Jensen, 2001; Schütz et al., 2001; Campler et al., 2009; Agnvall et al., 2012; Lindqvist and Jensen, 2009; Roth & Lind, 2013; Bessa Ferreira et al., 2022) as well as insights into how domestication and breed development can affect brain anatomy.

Here, we address this knowledge gap by quantifying the volumes of eight telencephalic regions of junglefowl and white leghorn chickens from the same populations used by Henriksen et al. (2016). In addition, we compared the relative sizes of telencephalic regions of chickens and junglefowl with four grouse species collected in the wild: ruffed grouse (*Bonasa umbellus*), sharp-tailed grouse (*Tympanuchus phasianellus*), lesser prairie-chicken (*Tympanuchus pallidicinctus*), and greater prairie-chicken (*Tympanuchus cupido*). These species were selected because we had an adequate number of male specimens in our brain collection, they were collected in the wild, and they are similar in body size to chickens and junglefowl. Based on previous work (Kruska and Schott, 1977; Ebinger and Löhmer, 1984, 1987; Kruska, 1988; Rehkämper et al., 1988; Ebinger, 1995; Ebinger and Röhrs, 1995) and the functional organization of the avian telencephalon (Shanahan et al., 2013), we predict several volumetric differences across species/strains and brain regions. First, limbic regions will be smaller in domestic chickens than in junglefowl or grouse because chickens are selected for reduced fear of humans (Agnvall et al., 2017), and a reduction in limbic regions is often found in comparisons of wild-domestic strains of mammals and birds ((Ebinger, 1995; Ebinger & Löhmer, 1984; Ebinger and Löhmer 1987; Ebinger and Röhrs, 1995; Kruska, 2005; Brusini et al., 2018). Second, chickens have lower visual acuity than junglefowl (Roth and Lind, 2013), so we expect the entopallium, the telencephalic target of the tectofugal visual pathway (Shimizu et al., 2010), to be reduced in domestic chickens compared to junglefowl. Third, we predict that all four grouse species will have larger hippocampus and sensory regions than chickens to support their navigational and sensory needs in the wild because chickens do not need to detect or avoid predators in captivity or search for food and shelter.

## Materials and Methods

### Specimens

To assess the effects of domestication on telencephalon composition, we measured the volumes of eight telencephalic regions in white leghorn chickens (n = 6), red junglefowl (n = 6) and four species of grouse: ruffed grouse (*Bonasa umbellus,* n = 6), greater prairie-chicken (*Tympanuchus cupido,* n = 3), lesser prairie-chicken (*Tympanuchus pallidicinctus,* n = 3), and sharp-tailed grouse (*Tympanuchus phasianellus,* n = 3). All specimens used were adult males, as determined by plumage and skull ossification (or known age in the case of the chickens and junglefowl). The chickens and junglefowl were obtained from breeding colonies maintained at Linköping University (Linköping, Sweden). The red junglefowl were derived from a Swedish zoo population and have been kept at Linköping University since 1998. The domesticated chickens originated from a selection line, SLU13, bred at the Swedish University of Agricultural Sciences. The sharp-tailed grouse specimens were obtained from hunters in Alberta, Canada. The ruffed grouse were trapped and euthanized in the field during the spring breeding season, as described in Krilow and Iwaniuk (2015). Both prairie-chicken species were live trapped on lekking grounds in Kansas and euthanized with cervical dislocation (Kansas Scientific Wildlife Permit SC-008-2014, Ohio State University IACUC permit 2013A00000013). All of these procedures adhered to the Guidelines for the Use of Wild Birds in Research (Fair et al. 2023).

For all specimens, the heads were removed as soon as possible after death and immersion fixed for several weeks in buffered 4% paraformaldehyde. The brains were extracted, weighed, cryoprotected, embedded in gelatin, and sectioned on a freezing stage microtome in the coronal plane at 40 µm. Every second section was mounted onto gelatinized slides and stained with thionin acetate.

### Quantification of Telencephalon Regions

Total telencephalon volume as well as the volumes of eight telencephalic regions were measured: hyperpallium, mesopallium, nidopallium, entopallium, arcopallium, septum, hippocampal formation (hippocampus proper plus area parahippocampalis), and striatum. The volumes were quantified using unbiased stereology with the Cavalieri estimator, as implemented in StereoInvestigator (Microbrightfield, Williston, VT), using a Zeiss Axio Imager M2 microscope (Carl Zeiss, MicroImaging GmBH, Germany). We referred to the chick (Puelles et al., 2018) and pigeon brain atlases (Karten & Hodos, 1967) to aid in defining borders for all regions measured. For hyperpallium volume, we took the combined volumes of both the dorsal and ventral hyperpallium as the two regions could not be consistently differentiated within and across specimens. Similarly, nucleus basalis could not always be discerned due to variation in staining quality, so we included nucleus basalis as part of our nidopallium measurements. To estimate the volumes of each brain region, a 1x lens was used and a grid size of either 400 µm x 400 µm or 450 µm x 450 µm (larger grid was used for the prairie-chickens and sharp-tailed grouse). Every 16^th^ section throughout the rostro-caudal extent of the telencephalon was measured for every specimen. Coefficients of error were all below 0.05 and the average data for chickens, junglefowl and each of the four grouse species are provided in Table 1.

**Table 1.**
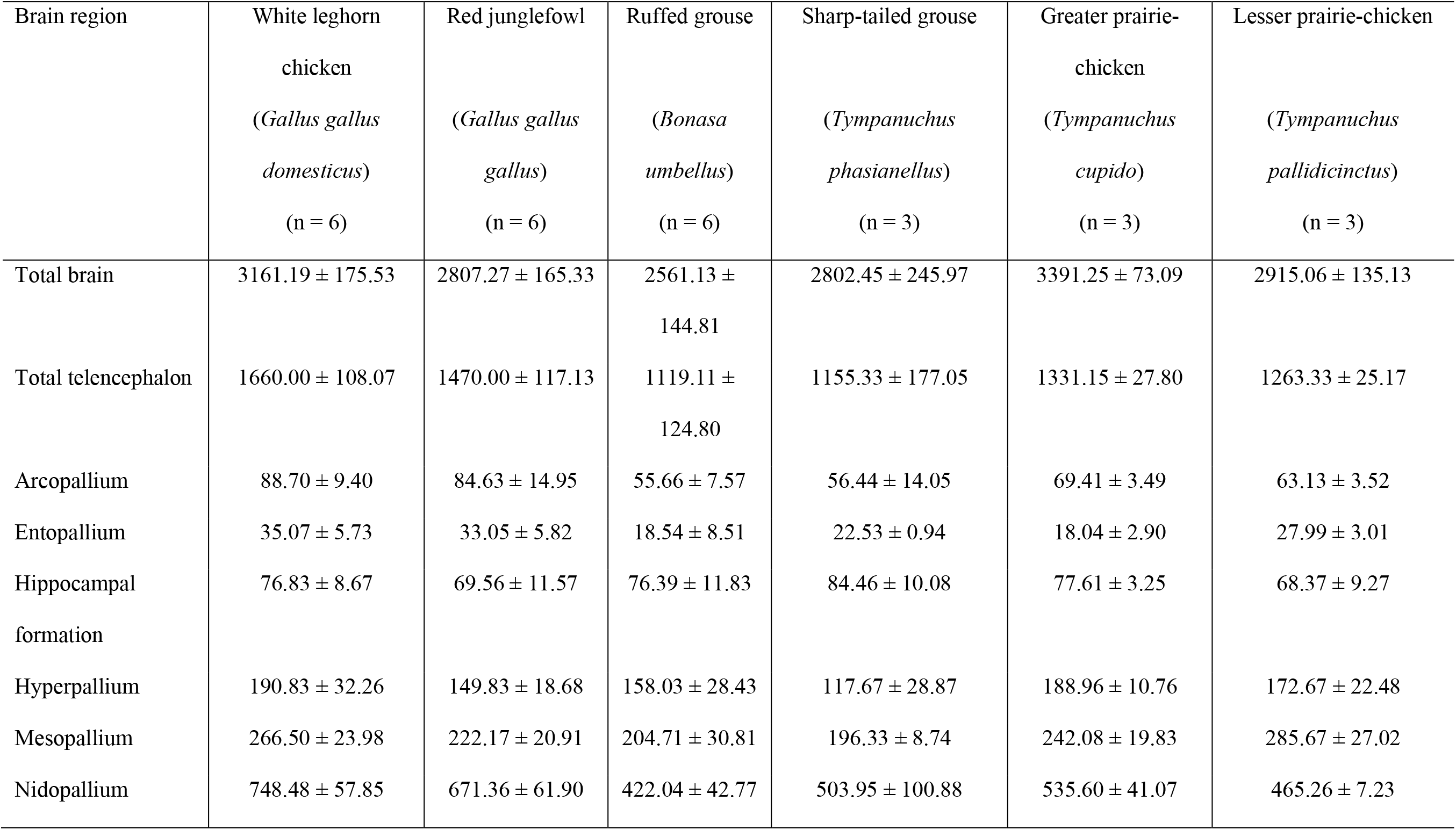
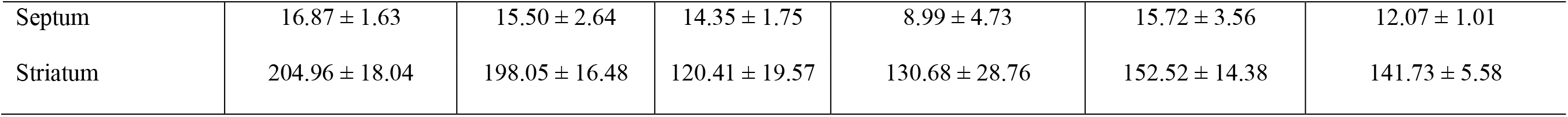
Volumetric measurements of the entire telencephalon and the eight telencephalic regions measured for this study for white leghorn chickens, red junglefowl and all four grouse species sampled. All measurements provided in mm^3^ as means ± standard deviations.

### Statistical Analyses

All data were log-transformed prior to statistical analysis. We first used one-way analyses of covariance (ANCOVAs) to test for significant differences in relative telencephalon region size among chickens, junglefowl, and ruffed grouse. We limited the ANCOVAs to these three because we had larger sample sizes (n = 6 for each) than for the remaining grouse species. Species/strain was used as a fixed factor, the telencephalon region of interest was used as the dependent variable (y-axis), and the telencephalon minus the region of interest was used as the covariate (x-axis). An interaction term was included initially for these comparisons to test for differences in allometric slopes among the three groups (see below). When significant differences were detected in our ANCOVA, we used Tukey’s post hoc tests to identify what pair-wise differences were present.

Because the telencephalon is heterogenous and different brain regions could be expanding/reducing in different species (or strains), we also used multivariate analyses. Specifically, we used both principal component analysis (PCA) and cluster analysis in a similar fashion to previous studies of brain composition across species (Iwaniuk and Hurd 2005) and chicken breeds (Rehkämper et al. 2003). For both sets of analyses, telencephalic region volume was expressed as a proportion of total telencephalon volume (i.e., region volume divided by total telencephalon volume) for each specimen. The PCA of the proportional volumes allowed us to reduce the total number of variables such that a smaller number of principal components (PCs) could be plotted in bivariate space. Cluster analysis provides an alternative means of multivariate analysis that provides a visual representation of similarities and differences across all individuals in the data set (Iwaniuk and Hurd 2005). For the cluster analysis, we used Euclidian distance measures with the ward D.2 clustering method. Euclidian distance measures (or the squared Euclidian distance measures) are recommended when using continuous variables (Yim & Ramdeen, 2015). The ward D.2 method creates clusters with the smallest variance between members of the same cluster, which allows for groups to form based on similarity to one another (Eszergár-Kiss & Caesar, 2017) and is therefore recommended for most datasets.

## Results

### Allometric comparisons of chickens, junglefowl, and ruffed grouse

No significant interaction effects between group (chicken, junglefowl, ruffed grouse) and the scaling variable (brain or telencephalon minus region of interest) were detected for our ANCOVAs (all p-values > 0.2), so we removed the interaction effects for all of the comparisons of relative telencephalon and telencephalon region volumes. Relative telencephalon size differed significantly among ruffed grouse, chickens and junglefowl (Table 2, Figure 1a). Post-hoc tests revealed that ruffed grouse have relatively smaller telencephalon volumes than both chickens and junglefowl and that junglefowl have relatively smaller telencephala than chickens, corroborating the findings of Henriksen et al. (2016).

**Fig. 1.**
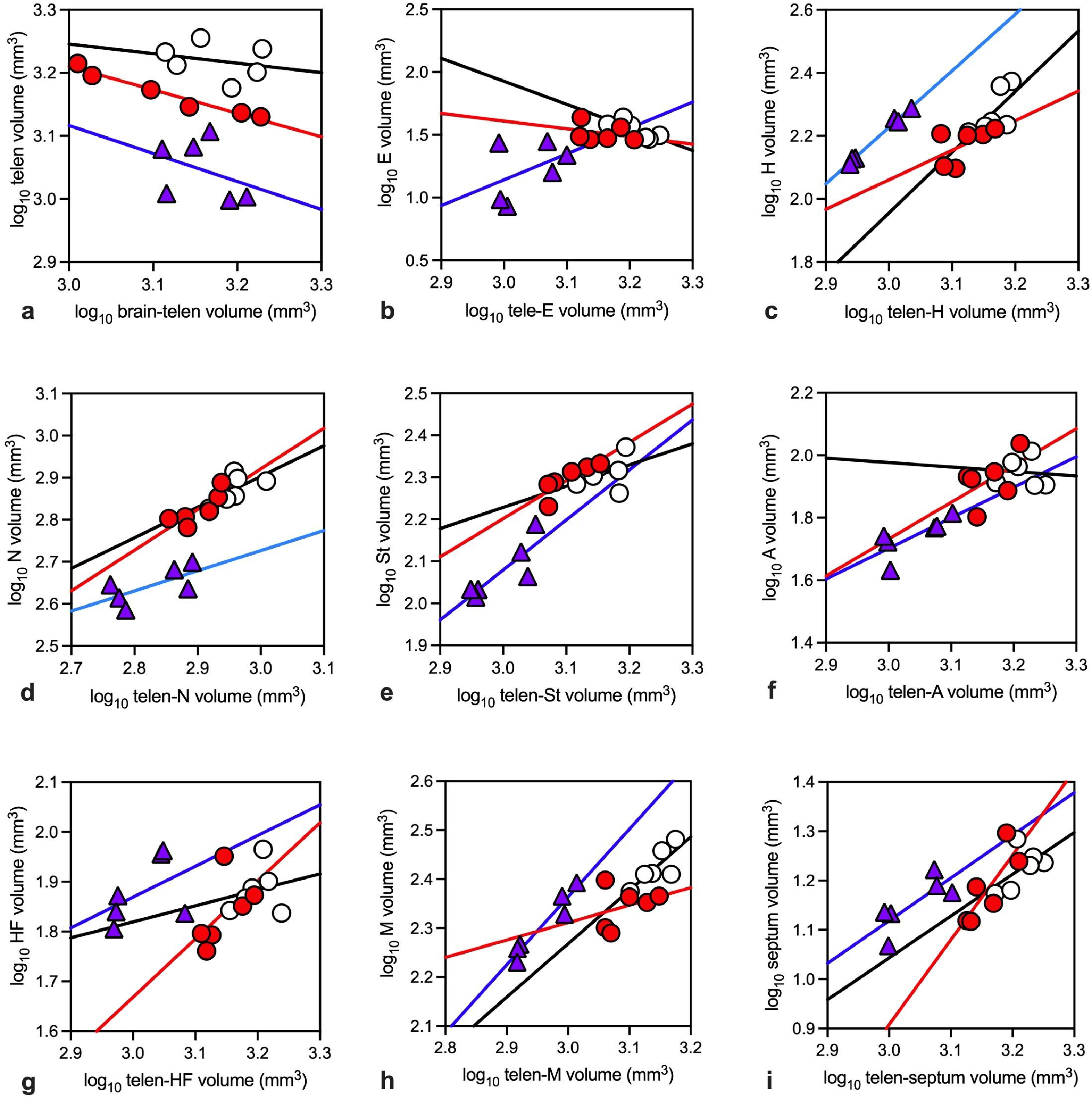
Scatterplots of brain region sizes relative to the rest of the brain (**a**) or rest of the telencephalon (**b-i**). The plots are as follows: telencephalon (**a**); entopallium (**b**); hyperpallium (**c**); nidopallium (**d**); striatum (**e**); arcopallium (**f**); hippocampal formation (**g**); mesopallium (**h**); and septum (**i**). In each plot, the blue triangles are ruffed grouse, the red circles are red junglefowl, and the white circles are white leghorn chickens. Least-squares linear regression lines are shown for each of these three: blue = ruffed grouse, red = red junglefowl, and black = white leghorn chicken. Abbreviated brain regions for the scatterplots are as follows: E = entopallium; H = hyperpallium; N = nidopallium; St = striatum; A = arcopallium; HF = hippocampal formation; and M = mesopallium.

**Table 2.**
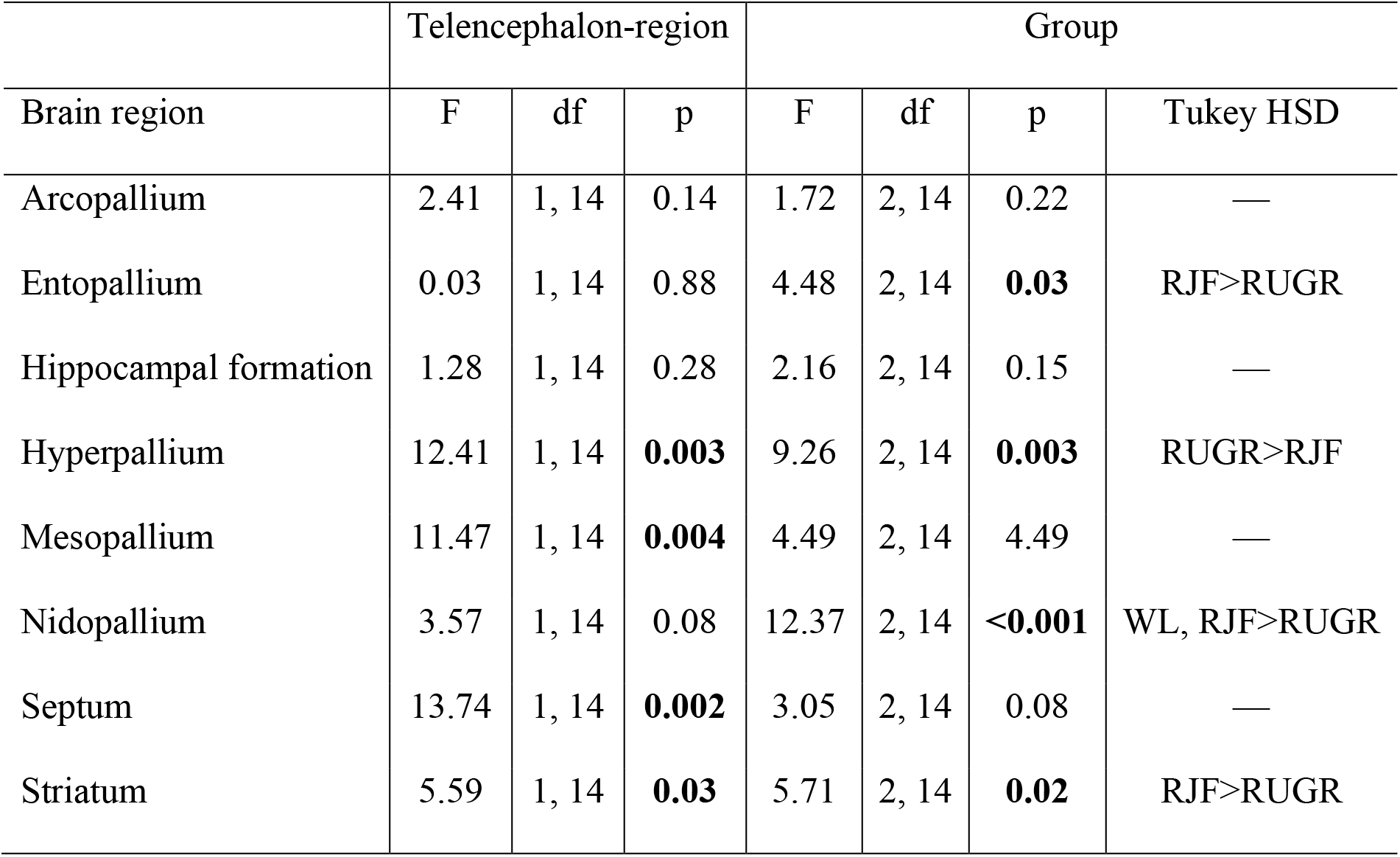
Results of analyses of covariance (ANCOVAs) of each of the eight telencephalic brain regions measured with telencephalon volume (minus region of interest) as a covariate and compared across red junglefowl (RJF), white leghorn chicken (WL) and ruffed grouse (RUGR) specimens. Significant effects are shown in bold. Where group was significant, the result of the post-hoc Tukey HSD tests are provided.

| Brain region | Telencephalon-region |  |  | Group |  |  |  |
| --- | --- | --- | --- | --- | --- | --- | --- |
|  | F | df | p | F | df | p | Tukey HSD |
| Arcopallium | 2.41 | 1, 14 | 0.14 | 1.72 | 2, 14 | 0.22 | — |
| Entopallium | 0.03 | 1, 14 | 0.88 | 4.48 | 2, 14 | **0.03** | RJF>RUGR |
| Hippocampal formation | 1.28 | 1, 14 | 0.28 | 2.16 | 2, 14 | 0.15 | — |
| Hyperpallium | 12.41 | 1, 14 | **0.003** | 9.26 | 2, 14 | **0.003** | RUGR>RJF |
| Mesopallium | 11.47 | 1, 14 | **0.004** | 4.49 | 2, 14 | 4.49 | — |
| Nidopallium | 3.57 | 1, 14 | 0.08 | 12.37 | 2, 14 | **<0.001** | WL, RJF>RUGR |
| Septum | 13.74 | 1, 14 | **0.002** | 3.05 | 2, 14 | 0.08 | — |
| Striatum | 5.59 | 1, 14 | **0.03** | 5.71 | 2, 14 | **0.02** | RJF>RUGR |

Across the eight telencephalic regions measured, we detected significant differences (Table 2) in the relative volumes of the entopallium (Figure 1b), hyperpallium (Figure 1c), nidopallium (Figure 1d), and striatum (Figure 1e). Based on post-hoc tests, the specific differences among the three groups varied across brain regions: the hyperpallium was relatively larger in ruffed grouse than red junglefowl, the nidopallium was relatively larger in both chickens and junglefowl than in ruffed grouse, and the entopallium and striatum were relatively larger in the junglefowl than the ruffed grouse. The remaining four brain regions, the arcopallium (Figure 1f), hippocampal formation (Figure 1g), mesopallium (Figure 1h), and septum (Figure 1i), did not differ significantly in relative size across groups (Table 2).

### Principal Component Analysis

Of the eight principal components, the first three comprise almost 76% of the variation across all individuals and species (Table 3). In the plot of principal component 2 (PC2) against PC1 (Fig. 2a), individuals are primarily separated along the PC1 axis (x-axis) with chickens and junglefowl on the left. Based on the loadings (Table 3, Fig. 2b), this indicates that the chickens and junglefowl have proportionally larger arcopallial, striatal and nidopallial volumes and smaller hippocampal, mesopallial, and hyperpallial volumes than the four grouse species. There was not, however, any apparent pattern or clustering along the PC2 axis, which largely reflects the proportional sizes of the entopallium, septum and mesopallium (Table 3, Fig. 2b). Plotting PC3 against PC1 again largely separates the chicken and junglefowl from the grouse along the PC1 axis (Fig. 2c). Along PC3, sharp-tailed grouse are slightly lower than many of the other data points, potentially indicating proportionally smaller hippocampal formation (Fig. 2d).

**Fig. 2.**
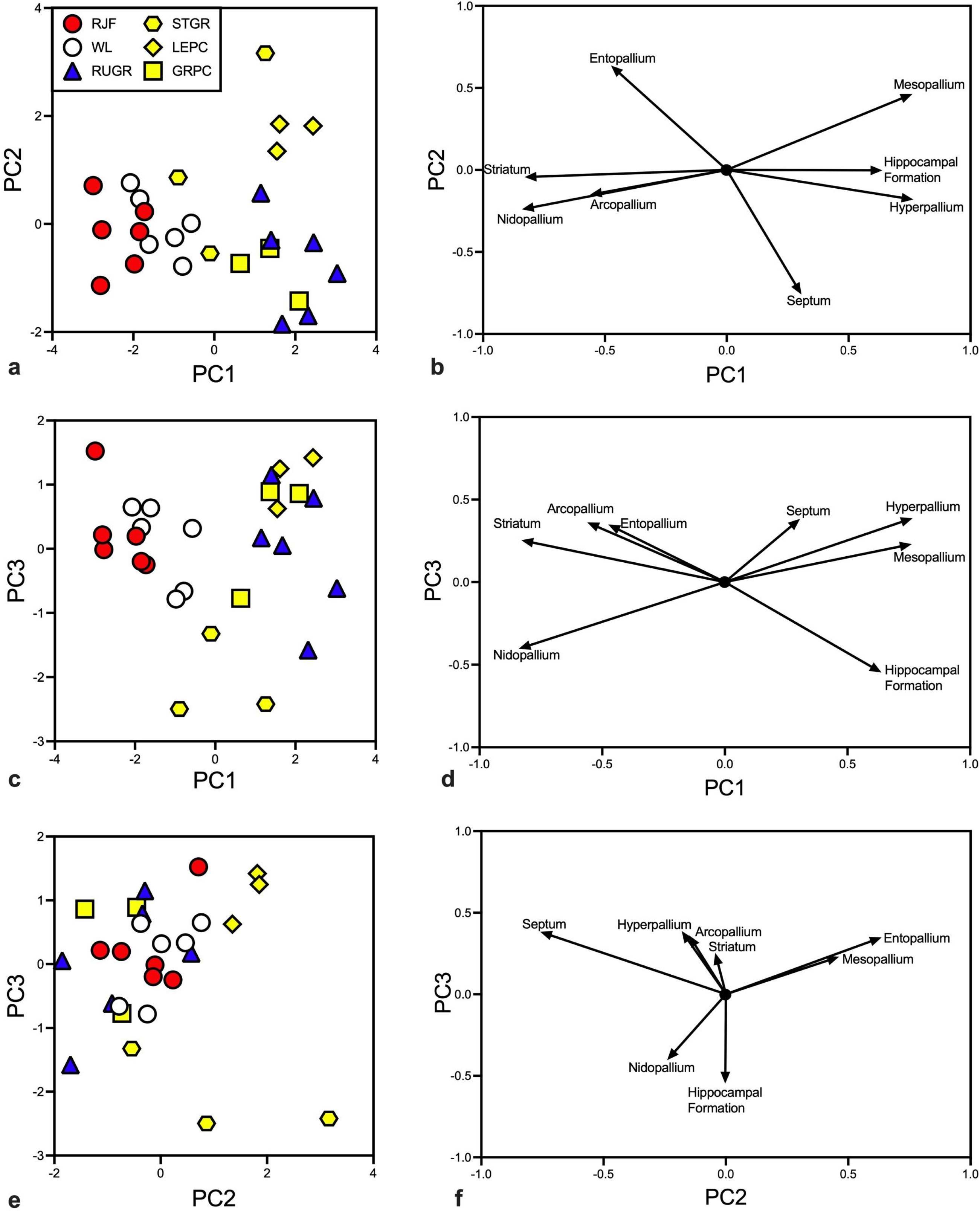
Scatterplots (**a, c, e**) and factor loadings of principal components (**b, d, f**) arising from a principal component analysis of the proportional sizes of eight telencephalic brain regions across white leghorn chickens (WL), red junglefowl (RJF), ruffed grouse (RUGR), sharp-tailed grouse (STGR), lesser prairie-chicken (LEPR), and greater prairie-chicken (GRPR). **a.** scatterplot of principal components (PC) 1 and 2 and a legend of the symbols shown in all three scatterplots. **b.** a loading plot of principal components 1 and 2. The arrows indicate the direction and magnitude of the loadings for each of the eight brain regions. **c.** scatterplot of principal components 1 and 3. **d.** a loading plot of principal components 1 and 3. **e.** scatterplot of principal components 2 and 3. **f.** a loading plot of principal components 2 and 3.

**Table 3.**
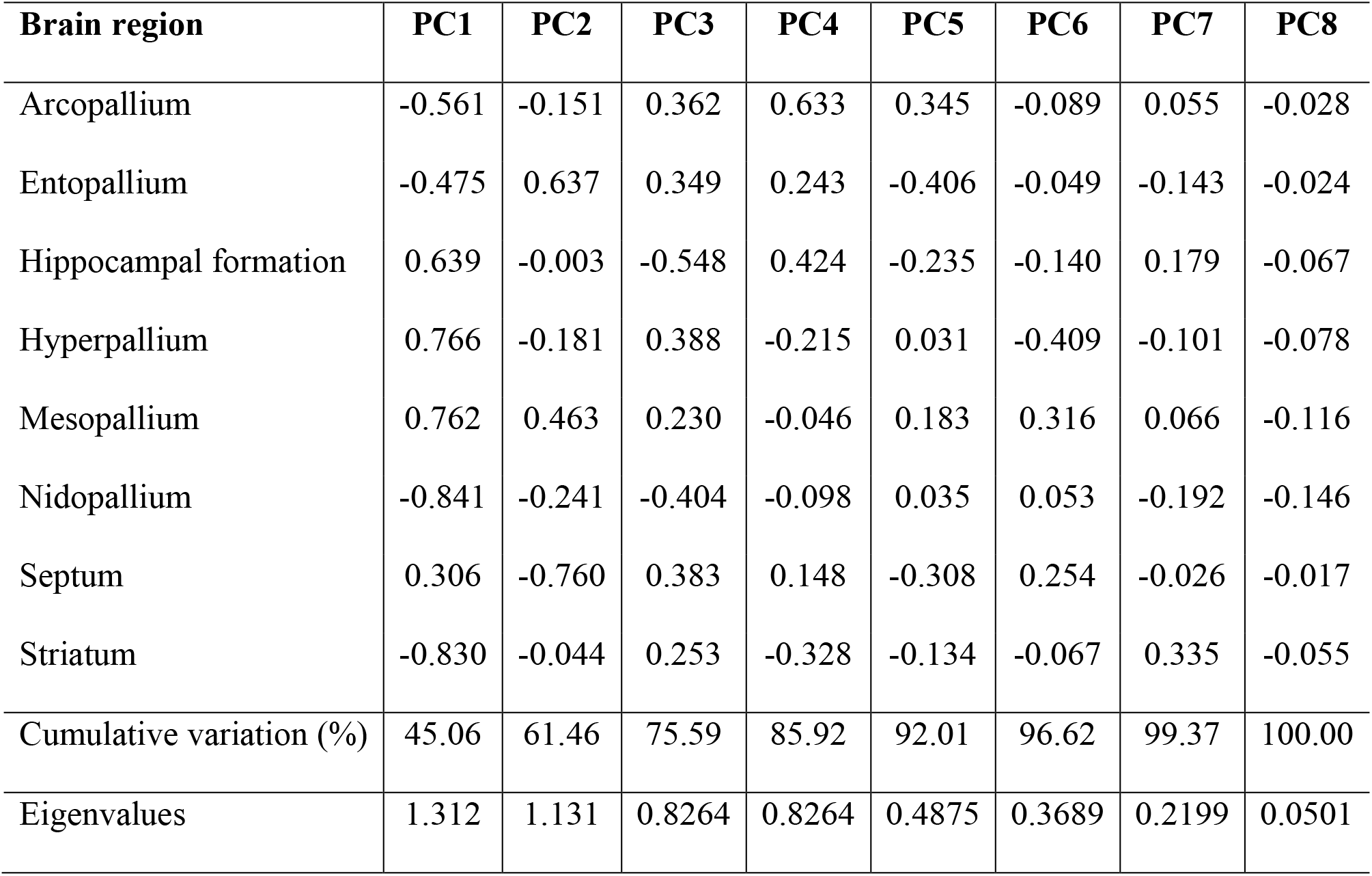
Loadings, cumulative variation, and eigenvalues for all eight principal components (PC) from the principal component analysis of the proportional sizes of eight telencephalic regions across chickens, junglefowl, and all four grouse species.

| **Brain region** | **PC1** | **PC2** | **PC3** | **PC4** | **PC5** | **PC6** | **PC7** | **PC8** |
| --- | --- | --- | --- | --- | --- | --- | --- | --- |
| Arcopallium | -0.561 | -0.151 | 0.362 | 0.633 | 0.345 | -0.089 | 0.055 | -0.028 |
| Entopallium | -0.475 | 0.637 | 0.349 | 0.243 | -0.406 | -0.049 | -0.143 | -0.024 |
| Hippocampal formation | 0.639 | -0.003 | -0.548 | 0.424 | -0.235 | -0.140 | 0.179 | -0.067 |
| Hyperpallium | 0.766 | -0.181 | 0.388 | -0.215 | 0.031 | -0.409 | -0.101 | -0.078 |
| Mesopallium | 0.762 | 0.463 | 0.230 | -0.046 | 0.183 | 0.316 | 0.066 | -0.116 |
| Nidopallium | -0.841 | -0.241 | -0.404 | -0.098 | 0.035 | 0.053 | -0.192 | -0.146 |
| Septum | 0.306 | -0.760 | 0.383 | 0.148 | -0.308 | 0.254 | -0.026 | -0.017 |
| Striatum | -0.830 | -0.044 | 0.253 | -0.328 | -0.134 | -0.067 | 0.335 | -0.055 |
| Cumulative variation (%) | 45.06 | 61.46 | 75.59 | 85.92 | 92.01 | 96.62 | 99.37 | 100.00 |
| Eigenvalues | 1.312 | 1.131 | 0.8264 | 0.8264 | 0.4875 | 0.3689 | 0.2199 | 0.0501 |

The third plot of PC3 against PC2 does not appear to show much differentiation across species (Fig. 2e), but the sharp-tailed grouse tend to be lower on the PC3 axis, largely due to relatively smaller hippocampal formation volumes (Fig. 2f).

### Cluster Analysis

The cluster analysis produced a dendrogram with two main groupings we labelled ‘A’ and ‘B’ (Fig. 3). Group ‘A’ is composed of the ruffed grouse, lesser prairie-chickens, and greater prairie-chickens and is characterized by larger hyperpallial, mesopallial, hippocampal, and nidopallial volumes, but smaller striatal, entopallial, and arcopallial volumes (Table 4). Group ‘B’ is composed of the red junglefowl, white leghorns, sharp-tailed grouse, and one greater prairie-chicken. This cluster is characterized by smaller hyperpallial, mesopallial, and nidopallial volumes, but larger striatal, entopallial, and arcopallial volumes (Table 4). Thus, the chickens and junglefowl tend to be more similar to one another than they are to the other species and the sharp-tailed grouse tends to differ from the other grouse species examined.

**Fig. 3.**
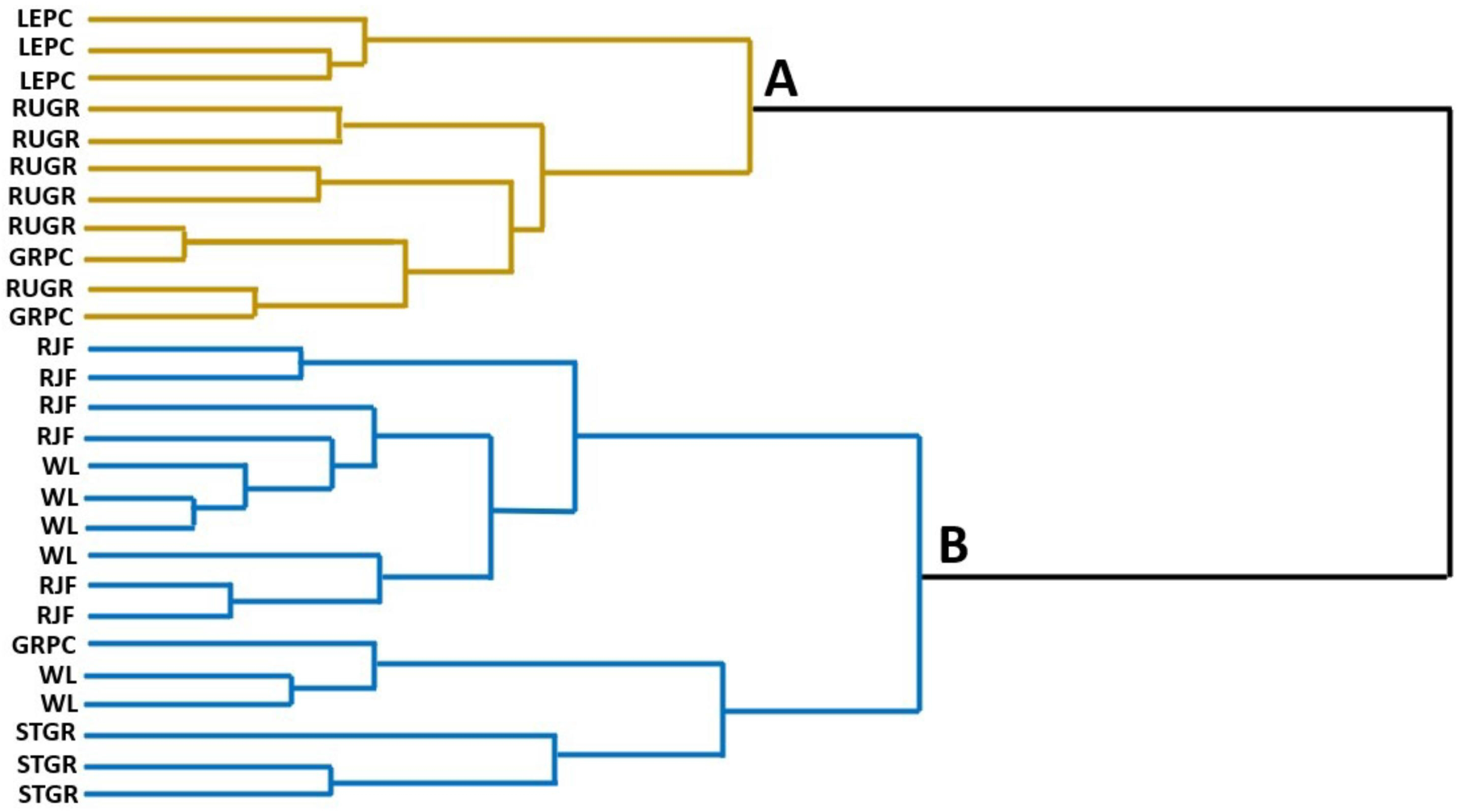
A dendrogram resulting from a UPGMA hierarchical cluster analysis. The two main clusters are indicated by the letters “A” (yellow, top) and “B “(blue, bottom). RJF = red junglefowl, WL = white leghorn chicken, RUGR = ruffed grouse, GRPC = greater prairie-chicken, LEPC = lesser prairie-chicken, STGR = sharp-tailed grouse. Cluster A is primarily composed of ruffed grouse and prairie-chickens. Cluster B is primarily composed of chickens, junglefowl and sharp-tailed grouse.

**Table 4.** The mean proportions of each of the eight telencephalic subregions in each cluster as created by UPGMA hierarchical cluster analysis. The brain regions are as follows: hyperpallium (H), mesopallium (M), nidopallium (N), striatum (St), entopallium (E), hippocampal formation (HF), septum, and arcopallium (A). The proportions were calculated by dividing the volume of each individual structure by that of the total telencephalon volume minus the subregion volume. The proportions for each cluster are the averages across every individual in that cluster. As shown in Figure 3, Cluster A is composed of ruffed grouse, lesser prairie-chicken, and greater prairie-chicken whereas Cluster B is composed of red junglefowl, white leghorn, sharp-tailed grouse, and one greater-prairie chicken. SD = standard deviation.

| Cluster |  | H | M | N | St | E | HF | Septum | A |
| --- | --- | --- | --- | --- | --- | --- | --- | --- | --- |
| A | mean | 0.1274 | 0.1901 | 0.4008 | 0.1116 | 0.0185 | 0.0628 | 0.0102 | 0.0508 |
|  | SD | ± 0.020 | ± 0.028 | ± 0.032 | ± 0.007 | ± 0.006 | ± 0.009 | ± 0.003 | ± 0.004 |
| B | mean | 0.1174 | 0.1634 | 0.4391 | 0.1239 | 0.0204 | 0.0525 | 0.0110 | 0.0536 |
|  | SD | ± 0.020 | ± 0.017 | ± 0.032 | ± 0.013 | ± 0.006 | ± 0.012 | ± 0.001 | ± 0.008 |

## Discussion

The relative size of the telencephalon differed between chickens and junglefowl, but not any of the telencephalic regions we measured. However, there were marked differences between both chickens and junglefowl in relative telencephalon size and the relative size of several telencephalic regions and the grouse species that we sampled. Thus, despite marked behavioural differences (Schütz and Jensen, 2001; Schütz et al., 2001; Campler et al., 2009; Agnvall et al., 2012; Lindqvist and Jensen, 2009; Roth & Lind, 2013; Bessa Ferreira et al., 2022), chickens and junglefowl are similar to one another in telencephalon composition and differ from that of several wild galliform species.

Before discussing the potential implications of these results, it is important to note two potential limitations of this study. First, we only examined males. Although this was advantageous for this study because our analyses were not confounded by varying numbers of males and females within each species or strain, there is potential for our results to differ when analyzing only females or integrating both males and females into a larger neuroanatomical study, particularly given how common sex differences in brain region sizes are across bird species (MacDougall-Shackleton & Ball, 1999; Nottebohm et al., 1976). Second, we used grouse as our “wild-type” group. Although grouse and chickens are both within the same family (Phasianidae), they are not each other’s closest relatives. Ideally, the wild species used in such a comparison would be the Chinese bamboo partridge (*Bambusicola thoracicus*) or francolins (Kimball et al., 2021), but obtaining these species in the wild is extremely difficult. The grouse represent wild specimens that are similar in body and brain size, even though they are not as closely related to the genus *Gallus* as other species (Kimball et al., 2021).

### Allometry of Telencephalon and Telencephalic Regions

Based on our analyses, chickens have relatively larger telencephalon volumes than junglefowl, which is consistent with the findings from Henriksen et al. (2016). In other domesticated birds, the telencephalon is relatively smaller in domesticates than wild strains: - 15% in ducks, −16% in geese, −24% in turkeys, and −7% in pigeons (Ebinger, 1995; Ebinger & Löhmer, 1984; Ebinger and Löhmer 1987; Ebinger and Röhrs, 1995). Thus, the chicken appears to be an outlier in that relative telencephalon size increases by approximately 17% rather than decreasing. One of the key differences between our study and previous works is that we used captive-bred junglefowl whereas other species comparisons used wild caught individuals.

Captivity can have effects on brain anatomy, even after as few as eight generations (Guay, 2008) and the junglefowl have been maintained in captivity far longer (Henriksen et al., 2021; Henriksen et al., 2016; Johnsson et al., 2018). However, it should be noted that true wild junglefowl are nearly impossible to source; almost all wild junglefowl populations have been hybridized with domesticated chickens (Peterson & Brisbin, 1998; Wu et al., 2020). So although an ideal comparison would be with wild red junglefowl, this no longer seems feasible and an unselected, captive bred population is the best option available.

Even if there is an effect of captivity on junglefowl brains, one would expect the reverse pattern based on behaviour and selection. Chickens are specifically selected for production: egg laying or meat. In the case of white leghorn chickens, this has been a particularly intense selection regime, and one would predict a relatively smaller brain and telencephalon due to allocating more resources to growth or egg production (van Schaik et al. 2023). Instead, chickens have proportionally enlarged the telencephalon compared with junglefowl (Fig. 1a) and have done so in a coordinated fashion such that the chicken telencephalon is a ‘scaled-up’ version of the red junglefowl. That is, chickens and junglefowl appear to share a similar telencephalic composition, differing only in the relative size of the telencephalon as a whole. This was unexpected given the differences in behaviour between the two strains (Schütz and Jensen, 2001; Schütz et al., 2001; Campler et al., 2009; Agnvall et al., 2012; Lindqvist and Jensen, 2009; Roth & Lind, 2013; Bessa Ferreira et al., 2022). However, there may be differences in neuron numbers, neuron density, or neurochemistry that would not be apparent in volumetric measurements. For example, nonapeptides (e.g., oxytocin, mesotocin, vasotocin) can affect many aspects of sociality and aggression (Bales et al., 2004; Carter et al., 2008; Goodson & Bass, 2001; Goodson et al., 2004a); infusions of mesotocin in zebra finch (*Taeniopygia guttata*) caused finches to become more social and spend more time in social groups (Goodson et al., 2009).

Given that chickens are much more tolerant of threats than junglefowl (Agnvall et al., 2014; Agnvall et al., 2012), it is possible that changes in neuropeptide levels or neuropeptide receptors are at least partially responsible for the behavioural differences between chickens and junglefowl. It is also possible that there are differences in neurotransmitters, particularly GABA and serotonin, as both seem to play a role in tameness (Albert et al., 2008) and several genes associated with neurotransmitters were identified as being involved in anxiety differences between chickens and junglefowl (Johnsson et al. 2016).

Unexpectedly, both chickens and junglefowl had significantly larger telencephala for their brain size than ruffed grouse. We predicted that ruffed grouse would have relatively larger telencephala to enable mate searching, predator avoidance, and finding food and other behaviours typically thought to be more frequent or “complex” in wild species. Our finding is not only the opposite of what we predicted, it is the opposite of what is typically observed in domesticate-wild comparisons (Ebinger, 1995; Ebinger & Löhmer, 1984; Ebinger and Löhmer 1987; Ebinger and Röhrs, 1995). How this might be related to behavioural differences can be potentially inferred from the volumetric differences identified in specific telencephalic regions.

Among the eight telencephalic regions measured, four of them differed significantly between ruffed grouse and junglefowl and/or chickens and four did not. The mixed results indicates that junglefowl and chickens have not enlarged all telencephalic regions equally, which was expected based on comparisons of domesticate-wild strains in other birds: some brain regions differed in size and others did not (Ebinger, 1995; Ebinger & Löhmer, 1984; Ebinger and Löhmer 1987; Ebinger and Röhrs, 1995). The arcopallium is one region that did not differ in relative size. As a premotor region (Shanahan et al. 2013), is it likely that it shares similar processing requirements across uncaged chickens, junglefowl and ruffed grouse. Similarly, the septum is a highly conserved brain region associated with memory, social behaviour and several other functions (Goodson 2005; Goodson et al. 2004b) that are shared across the three groups.

Further, behavioural variation within and across species is more typically associated with neurochemical changes in the septum (Goodson 2005; Goodson et al. 2004b). The remaining two regions that did not differ significantly, the hippocampal formation and mesopallium, were slightly larger in ruffed grouse than chicken and junglefowl, but these differences were not significant and a lot of inter-individual variation was found within each group.

Of the four regions that did differ significantly in relative size, only the hyperpallium was larger in ruffed grouse than chickens and junglefowl. The hyperpallium processes primarily somatosensory and visual information (Atoji et al., 2018). The functional relevance of a relatively larger or smaller hyperpallium is challenging to interpret given its multisensory nature (Reiner et al., 2005; Shanahan et al., 2013). It is possible that ruffed grouse differ in sensory abilities from chickens and junglefowl, but this would require behavioural testing.

The remaining three regions, entopallium, nidopallium, and striatum, were all larger in junglefowl and/or chickens than ruffed grouse. The difference in relative entopallium size is difficult to interpret because entopallium-telencephalon allometry was not significant and the ruffed grouse exhibited more inter-individual variation than the chickens and junglefowl (Fig. 1b). Therefore, this appears to be largely an absolute difference in entopallium size and it is unclear if this reflects visual system function. Conversely, the differences in relative nidopallium and striatum volumes appear to be more robust. The nidopallium and striatum comprise over 50% of telencephalic volume and their expansion is likely responsible for the differences in relative telencephalon size (Fig. 1a) and potentially behavioural differences. The nidopallium is the largest subregion within the telencephalon and has many functions and numerous subdivisions (Shanahan et al., 2013). In general, the nidopallium is responsible for higher-level processing, sensory integration, and cognition (Shanahan et al., 2013), suggesting differences in cognition and/or sensory processing that could be assessed using a battery of behavioural tests across species/strains (Cauchoix et al., 2017; Shaw, 2017). The striatum is part of the basal ganglia circuitry, and as such plays a significant role in movement control and coordination (Reiner et al., 1998; Reiner et al., 2005). This could reflect a difference in motor control or coordination between chickens and junglefowl and ruffed grouse, but again requires behavioural testing.

### Multivariate Analyses of Telencephalon Composition

The principal component analysis (PCA) revealed that chickens and junglefowl have a different telencephalon composition than other grouse species. To elaborate, junglefowl and chickens’ group together with sharp-tailed grouse, largely due to differences in hyperpallium, mesopallium, and nidopallium sizes compared to other grouse. With such a small number of species sampled, it is not possible to determine why the sharp-tailed grouse groups with the chickens and junglefowl and not with other members of its own genus (i.e., the prairie-chickens). However, the fact that the junglefowl and chickens group together in both the cluster analysis and PCA suggests that they share a different “cerebrotype” (Iwaniuk & Hurd, 2005) that is different from wild galliform species. The chicken and junglefowl may be more similar to one another because they are two strains of the same species, but this does not fully explain why both of them differ from other galliforms. Based on the loadings of the PCA and cluster analysis, the *Gallus* cerebrotype appears to primarily reflect relatively larger nidopallial and striatal volumes and smaller mesopallial and hyperpallia volumes. To some extent, this corroborates our comparisons with ruffed grouse (see above) and indicates that chickens and junglefowl differ in telencephalon composition from several wild galliforms and not just ruffed grouse. Once more, the implication is that this might reflect significant behavioural differences, but requires a battery of behavioural tests.

## Conclusions

Overall, chickens and junglefowl do not differ significantly from one another in telencephalic composition, but do differ from ruffed grouse and other grouse species. Although we cannot completely discount the effects of captive breeding, the enlargement of some brain regions in both chickens and junglefowl suggests that *Gallus gallus* differs in how the brain has changed in response to domestication from other domesticated species, such as pigeons, ducks, geese, and turkeys (Ebinger, 1995; Ebinger & Löhmer, 1984; Ebinger and Löhmer 1987; Ebinger and Röhrs, 1995). Our results also imply marked behavioural differences between chickens/junglefowl and other galliforms that should be tested experimentally in the lab and field, such as visual discrimination, motor coordination, and an assortment of cognitive tasks. Further analyses of chicken, junglefowl and other galliform species brains should also be conducted to determine more specifically what is different about chickens and junglefowl. This should include quantification of neuron numbers and neuron morphology across multiple telencephalic regions as well as expression of neurotransmitters, receptors, and other neurochemical markers associated with anxiety, stress, and neuroplasticity (Hecht et al. 2023).

## Acknowledgements

We thank Ben Brinkman and Maurice Needham for help with microscopy and Justin Krilow for assistance in obtaining both greater and lesser prairie-chickens. Funding was provided by scholarships to KJR from the University of Lethbridge and grants to ANI from the Natural Science and Engineering Research Council of Canada, Canada Foundation for Innovation, and Canada Research Chairs Program.

## Statement of ethics

All procedures complied with all relevant ethical regulations and were conducted under approvals and licenses in Canada, Sweden, and the USA.

## Conflict of Interest Statement

The authors have no conflicts of interest to declare.

## Funding Sources

All histology and analyses were supported by grants to A.N.I. from the Natural Sciences and Engineering Research Council of Canada and the Canada Research Chairs Program.

## Author Contributions

All authors had full access to all the data in the study and take responsibility for the integrity of the data and the accuracy of the data analysis. Study concept and design: K.J.R., R.H., and A.N.I. Acquisition of specimens: J.A., R.H., D.W., and A.N.I. Acquisition of the data: K.J.R. and J.R.H. Analysis and interpretation of data: K.J.R., R.H., and A.N.I. Drafting of the manuscript: K.J.R. and A.N.I. Critical revision of the manuscript for important intellectual content: K.J.R., J.A., R.H., D.W., and A.N.I. Obtained funding: A.N.I. Administrative, technical, and material support: J.A., R.H., D.W., and A.N.I. Study supervision: A.N.I.

## Notes

### Competing Interest Statement

The authors have declared no competing interest.

